# Deleting *Mecp2* from the entire cerebellum rather than its neuronal subtypes causes a delay in motor learning in mice

**DOI:** 10.1101/2020.11.12.380162

**Authors:** Nathan P. Achilly, Lingjie He, Olivia A. Kim, Shogo Ohmae, Gregory J. Wojaczynski, Tao Lin, Roy V. Sillitoe, Javier F. Medina, Huda Y. Zoghbi

## Abstract

Rett syndrome is a devastating childhood neurological disorder caused by mutations in *MECP2*. Of the many symptoms, motor deterioration is a significant problem for patients. In mice, deleting *Mecp2* from the cortex or basal ganglia causes motor dysfunction, hypoactivity, and tremor, which are abnormalities observed in patients. However, little is known about the consequences of deleting *Mecp2* from the cerebellum, a brain region critical for motor function. Here we show that deleting *Mecp2* from the entire cerebellum, but not from individual cerebellar cell types, causes a delay in motor learning that is overcome by additional training. We also observed irregular firing rates of Purkinje cells and transcriptional misregulation within the cerebellum of knockout mice. These findings demonstrate that the motor deficits present in Rett syndrome arise, in part, from cerebellar dysfunction. For Rett syndrome as well as other neurodevelopmental disorders, our results highlight the importance of understanding which brain regions contribute to disease phenotypes.

## INTRODUCTION

Loss-of-function mutations in *MECP2* cause a severe childhood disorder called Rett syndrome (Amir et al., 1999). After a period of normal development, patients lose previously acquired milestones and develop debilitating neurological deficits (Hagberg, et al., 1983; Neul et al., 2014). Of these symptoms, motor deterioration is a significant problem for patients and manifests as ataxia, apraxia, hypotonia, and spasticity (Neul et al., 2014; Sandweiss et al., 2020). In mice, the complete loss of *Mecp2* causes motor deficits such as ataxia, hind limb clasping, hypoactivity, and tremor that mimic those seen in patients (Chen et al., 2001; Guy et al., 2001). Furthermore, mice that conditionally lack *Mecp2* in the cortex or basal ganglia partially replicate these Rett-like impairments (Chen et al., 2001; Gemelli et al., 2006; Su et al., 2015), suggesting that forebrain dysfunction contributes to the motor deficits seen in patients. However, other brain regions such as the cerebellum also control motor activity (Bostan and Strick, 2019), and thus may also contribute to the complex, wide-ranging motor phenotypes of Rett syndrome.

The cerebellum contains approximately 75% of all neurons in the brain (Lang et al., 1975; Sarko et al., 2009) and integrates sensory inputs in order to fine-tune motor output (Manto et al., 2012). This function is critical for motor coordination and motor learning as impairments in cerebellar circuity cause ataxia, dystonia, and tremor (White et al., 2016; Bostan and Strick, 2019; Darmohray et al., 2019; Machado et al., 2020). The cerebellum also contributes to non-motor behaviors such as social interaction, reward, and memory (Wagner et al., 2017; Carta et al., 2019; McAfee et al., 2019; Kelly et al., 2020). Interestingly, some of the motor and non-motor symptoms of Rett syndrome overlap with those of conditions that perturb cerebellar function such as spinocerebellar ataxias, tumors, and strokes (Schmahmann, 2004; Sokolov, 2018; Sandweiss et al., 2020). Therefore, we hypothesized that MeCP2 deficiency disrupts cerebellar function leading to motor incoordination phenotypes similar to those seen in Rett syndrome. To test this hypothesis, we deleted *Mecp2* from the cerebellum and discovered that cerebellar knockout animals had deficits in motor learning that, interestingly, were overcome with additional training. This motor learning delay was accompanied by abnormal firing properties of Purkinje cells and an altered transcriptional profile of cerebellar neurons. These data indicate that MeCP2 deficiency in the cerebellum is consequential and contributes to the motor dysfunction seen in Rett syndrome.

## RESULTS

### Deleting *Mecp2* from the entire cerebellum causes a delay in motor learning

To confirm that MeCP2 is expressed in cerebellar neurons, we performed immunostaining for MeCP2 and a variety of neuron-specific makers in adult wild-type mice. MeCP2 was expressed in granule cells, Purkinje cells, and interneurons (Figure 1A-D). This suggests that MeCP2 may contribute to the function of these neuronal populations.

**Figure 1.**
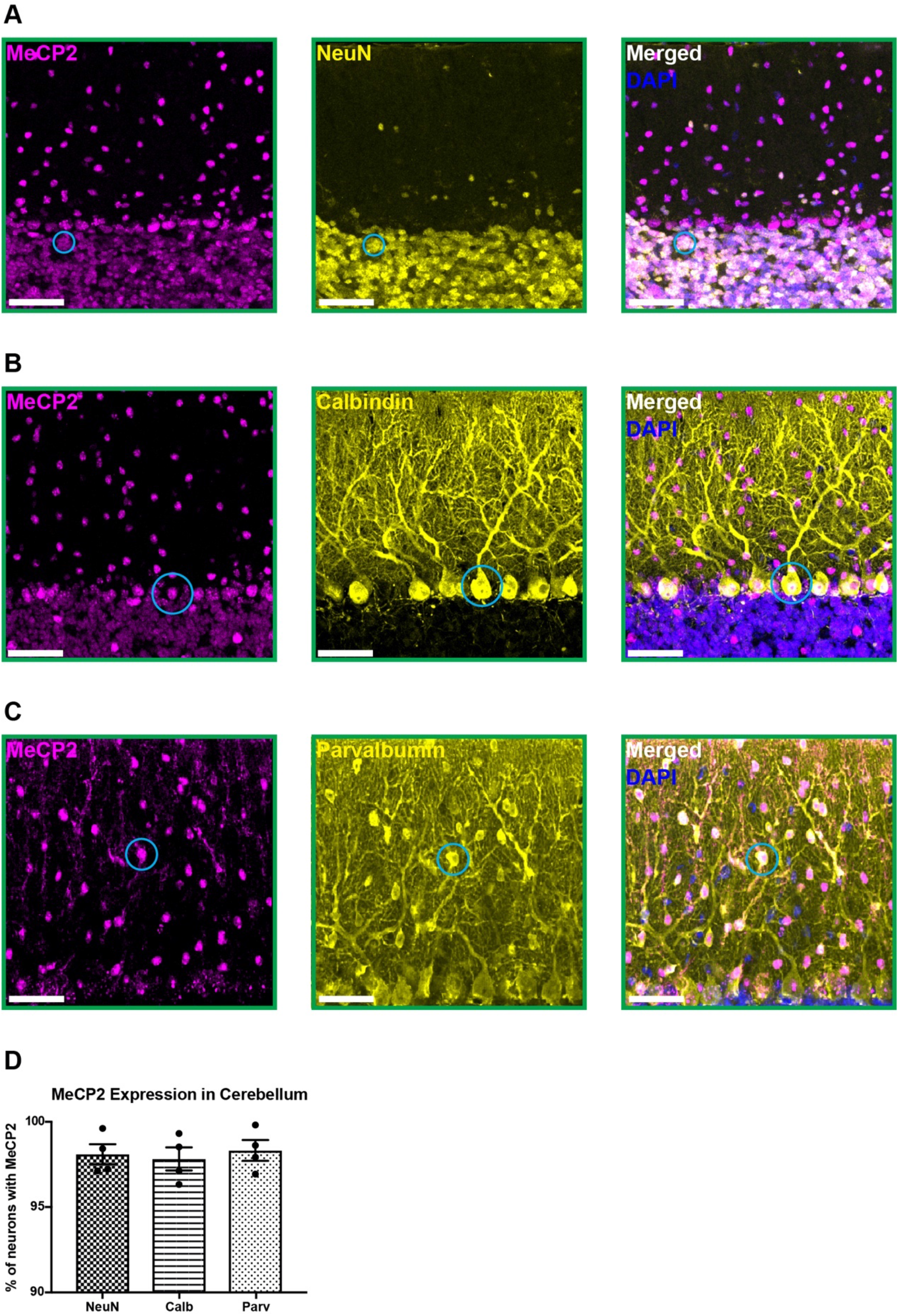
MeCP2 is expressed in cerebellar neurons of wild-type mice. (**A-C**) MeCP2 (magenta) staining in NeuN+ neurons (yellow) in the granular layer (A: solid cyan circle), Calbindin+ neurons (yellow) in the Purkinje cell layer (**B**: solid cyan circle), and Parvalbumin+ neurons (yellow) in the molecular layer (**C**: solid cyan circle). Scale bar, 25 μm. (**D**) Quantification of the percentage of NeuN+, Calbindin+, and Parvalbumin+ neurons that express MeCP2. N = 4 biologically independent mice per group. Data are presented as mean ± s.e.m.

To test this, we conditionally deleted *Mecp2* either from the entire cerebellum using *En1-Cre* mice, or individually from granule cells, Purkinje cells, and interneurons using *Atoh1-Cre*, *L7-Cre*, and *Ptf1a-Cre* mice, respectively (Hoshino et al., 2005; Joyner and Zervas, 2006; Hashimoto and Hibi, 2012; Slugocka et al., 2017). We verified the recombination efficiency using *Rosa26-tdTomato* reporter mice and found that granule cells, Purkinje cells, and interneurons expressed tdTomato (Figure 2-figure supplement 1A-C; Figure 2-figure supplement 2A-B,D-E,G-H). We crossed *Cre*-expressing and *Mecp2*^flox/+^ animals to generate *Mecp2* conditional knockout (*KO*) mice and littermate controls and confirmed that MeCP2 was deleted from the entire cerebellum or from individual cell types (Figure 2-figure supplement 1D-E; Figure 2-figure supplement 2C,F,I).

Because the cerebellum is involved in motor coordination and learning (Mauk et al., 2000), we analyzed the motor performance of cerebellar *KO* mice using the rotarod (Deacon, 2013). In this assay, mice are placed on a rotating rod that gradually accelerates. After four trial days, healthy mice spend progressively more time on apparatus as their motor skill improves, while mice with motor learning impairments spend less time on the apparatus. Although the performance of two- and four-month-old cerebellar *KO* mice was normal compared to control mice, six-month-old cerebellar *KO* mice had an initial motor learning delay that was overcome with additional training (Figure 2A; Figure 2-figure supplement 3A-B). These motor learning deficits were not due to abnormalities in general locomotor activity or strength (Figure 2-figure supplement 3C-F). Although the cerebellum is implicated in non-motor behaviors (Klein et al., 2016), we did not observe any deficits in sensorimotor gating, social behavior, or contextual fear memory in cerebellar *KO* mice (Figure 2-figure supplement 3G-K).

**Figure 2.**
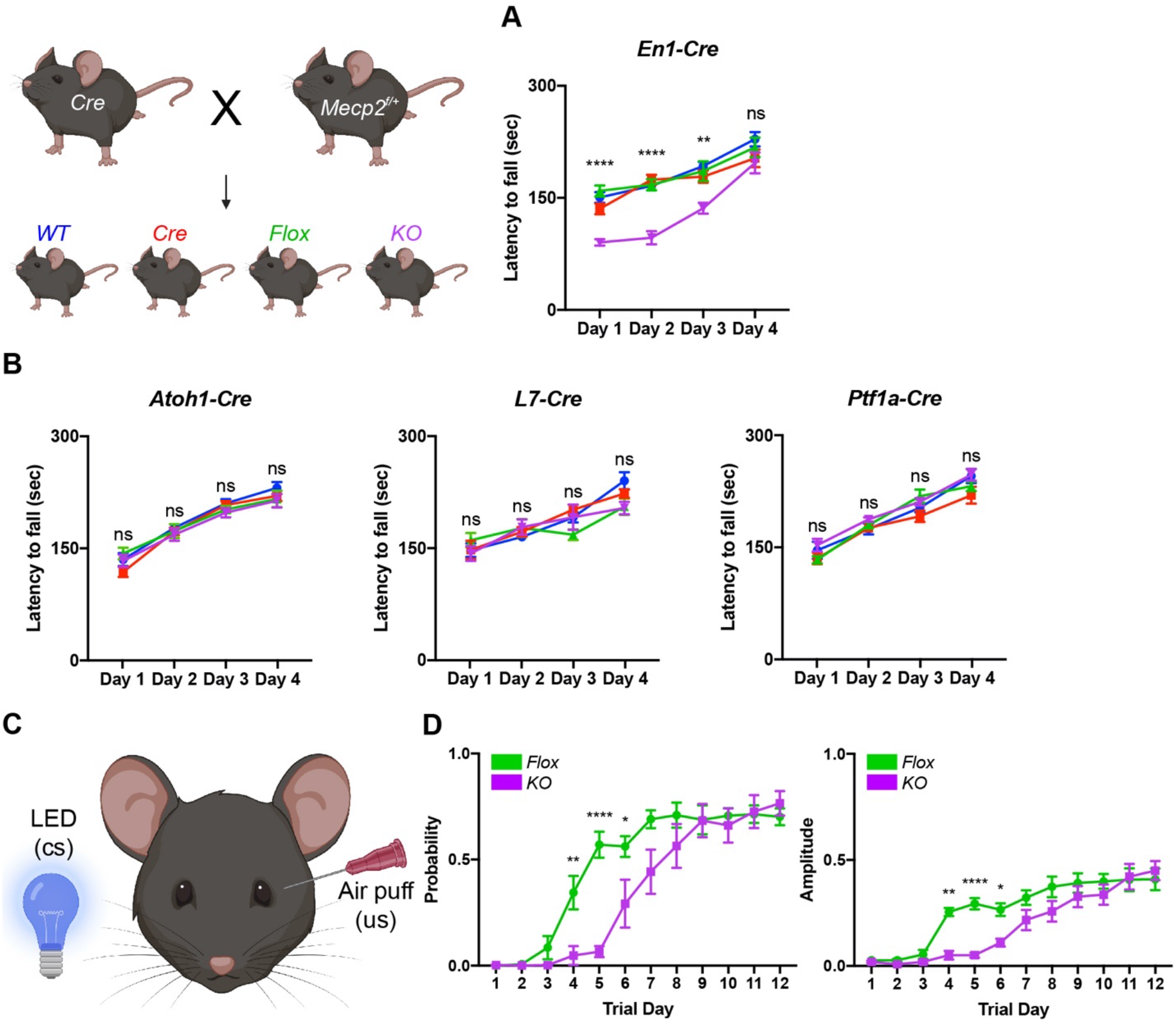
Deleting *Mecp2* from the entire cerebellum but not individual cell types causes motor learning deficits in 6-month-old mice. (**A**) Latency to fall on the rotarod over four training days in the *En1-Cre* group. (**B**) Latency to fall on the rotarod over four training days in mice lacking *Mecp2* in the granule cells (*Atoh1-Cre*), Purkinje cells (*L7-Cr*e), and interneurons (*Ptf1a-Cre*). (**C**) Schematic of eyeblink conditioning that pairs an LED light (conditioned stimulus, cs) with an air puff (unconditioned stimulus, us) to generate an anticipatory eyelid closure (conditioned response) before the air puff. (**D**) Response probability and amplitude of eyelid closure over 12 training days in *Flox* and *KO* mice. N = 8-17 biologically independent mice per group. Data are presented as mean ± s.e.m. ns(p>0.05), *(p<0.05), **(p<0.01), ****(p<0.0001).

Studies from other *Mecp2* knockout mice have demonstrated that behavioral deficits originate from dysfunction in a particular cell-type (Chao et al., 2010; Ito-Ishida et al., 2015; Meng et al., 2016; Mossner et al. 2019). For example, altered social behavior and seizures in *Mecp2* knockout mice originate from dysfunction in parvalbumin-expressing and somatostatin-expressing inhibitory neurons, respectively (Chao et al. 2010; Ito-Ishida et al. 2015). Therefore, we hypothesized that the motor learning deficits in cerebellar *KO* mice originate from the loss of MeCP2 in Purkinje cells, granule cells, or interneurons. However, we did not detect rotarod deficits in cell type-specific knockout animals at an age when cerebellar *KO* mice were symptomatic (Figure 2B).

In addition to the cerebellum, multiple brain regions including the cortex and basal ganglia contribute to motor learning on the rotarod (Sakayori et al., 2019). Therefore, we used eyeblink conditioning, an alternate task of motor learning that is entirely driven by the cerebellum, to validate the motor learning deficits observed in cerebellar *KO* mice (Figure 2C) (Heiney et al., 2014). Compared to control mice, cerebellar *KO* mice also exhibited an initial delay in the probability and amplitude of eyelid closure that improved with additional training, suggesting that cerebellar-dependent motor learning is disrupted by the loss of *Mecp2* (Figure 2D). These results demonstrate that deleting *Mecp2* from the cerebellum causes motor learning delay.

### Purkinje cell firing rate is more variable in cerebellar knockout mice

Because removing *Mecp2* perturbs synaptic function (Na et al., 2013; Meng et al., 2016; Ure et al., 2016), we hypothesized that alterations in the electrophysiological properties of cerebellar neurons might explain the motor phenotypes in cerebellar *KO* mice. To accomplish this, we monitored the activity of Purkinje cells in awake animals by recording their simple spikes, which are modulated by granule cells, and complex spikes, which are triggered by input from inferior olivary neurons (Schmolesky et al., 2002; Davie et al., 2008; Arancillo et al., 2015) (Figure 3A-C). Although the mean firing rates were unchanged, the simple spike coefficient of variation 2 (CV2), a measure of firing rate irregularity, was elevated in cerebellar *KO* mice (Figure 3D). This electrophysiological abnormality was not due to defects in Purkinje cell morphology or the density of excitatory and inhibitory synaptic puncta (Figure 3E-F). Thus, deleting *Mecp2* from the cerebellum disrupts some aspects of Purkinje cell firing in absence of overt morphological abnormalities.

**Figure 3.**
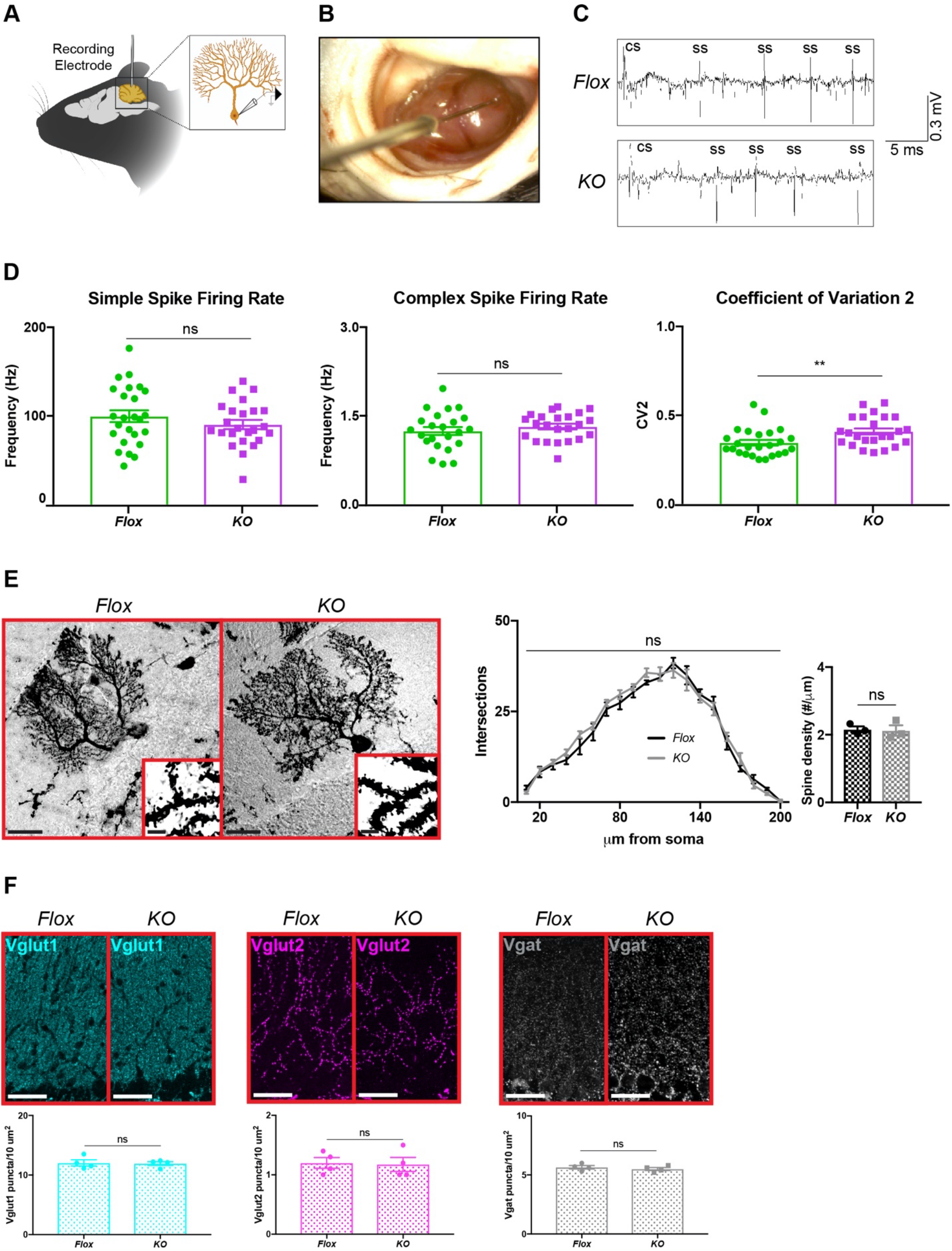
Purkinje cell firing rate is more variable in *KO* mice but is independent of overt morphological abnormalities. (**A**) Schematic of in vivo extracellular recording of Purkinje cells. (**B**) Photograph of a recording electrode inside a surgically-implanted recording chamber. (**C**) Representative traces of Purkinje cell firing pattern in *Flox* and *cKO* mice displaying simple spikes (ss) and complex spikes (cs). (**D**) Simple spike firing rate, complex spike firing rate, and coefficient of variation 2 (CV2). (**E**) Golgi stain of cerebellar Purkinje cells in *Flox* and *KO* mice. Scale bar, 25 μm. Inner panel demonstrates dendritic spines on Purkinje cells. Scale bar, 5 μm. Sholl analysis and spine density quantification in *Flox* and *KO* mice. (**F**) Staining and quantification of Vglut1 (cyan), Vglut2 (magenta), and Vgat (gray) puncta density in the cerebellum of *Flox* and *KO* mice. Scale bar, 25 μm. For (**D**), N = 23-27 neurons from 3 biologically independent mice per group. For (**G**), 10-15 neurons were analyzed from 3 biologically independent mice per group. For (**H**), N = 4 biologically independent mice per group. Data are presented as mean ± s.e.m. ns(p>0.05), **(p<0.01).

### Transcriptional misregulation occurs in cerebellar knockout mice

MeCP2 is a transcriptional regulator, and gene misregulation is a hallmark of its dysfunction (Ben-Shachar et al., 2009; Zhao et al., 2013; Johnson et al., 2017; Sanfeliu et al., 2019). In the cerebellum of global *Mecp2* null mice, approximately 100 genes are misregulated by more than 20% relative to wild-type mice (Ben-Shachar et al., 2009; Raman et al., 2018). These gene expression abnormalities could result from the absence of MeCP2 from the cerebellum or from the absence of MeCP2 from the surrounding brain. To determine if transcriptional misregulation occurs in cerebellar *KO* mice and if it is caused by cell-autonomous or non-cell-autonomous factors, we performed RT-qPCR for 20 genes whose expression is most altered in the cerebellum of global *Mecp2* null mice (Raman et al., 2018). The levels of *Egr4, Cabp7, Rreb1*, and *Vav3* were altered in six-month-old, symptomatic *KO* mice but not in two-month-old, asymptomatic *KO* mice (Figure 4A-B). Thus, deleting *Mecp2* from the cerebellum mimics some gene expression changes in global *Mecp2* null mice, and the degree of misregulation worsens with age.

**Figure 4.**
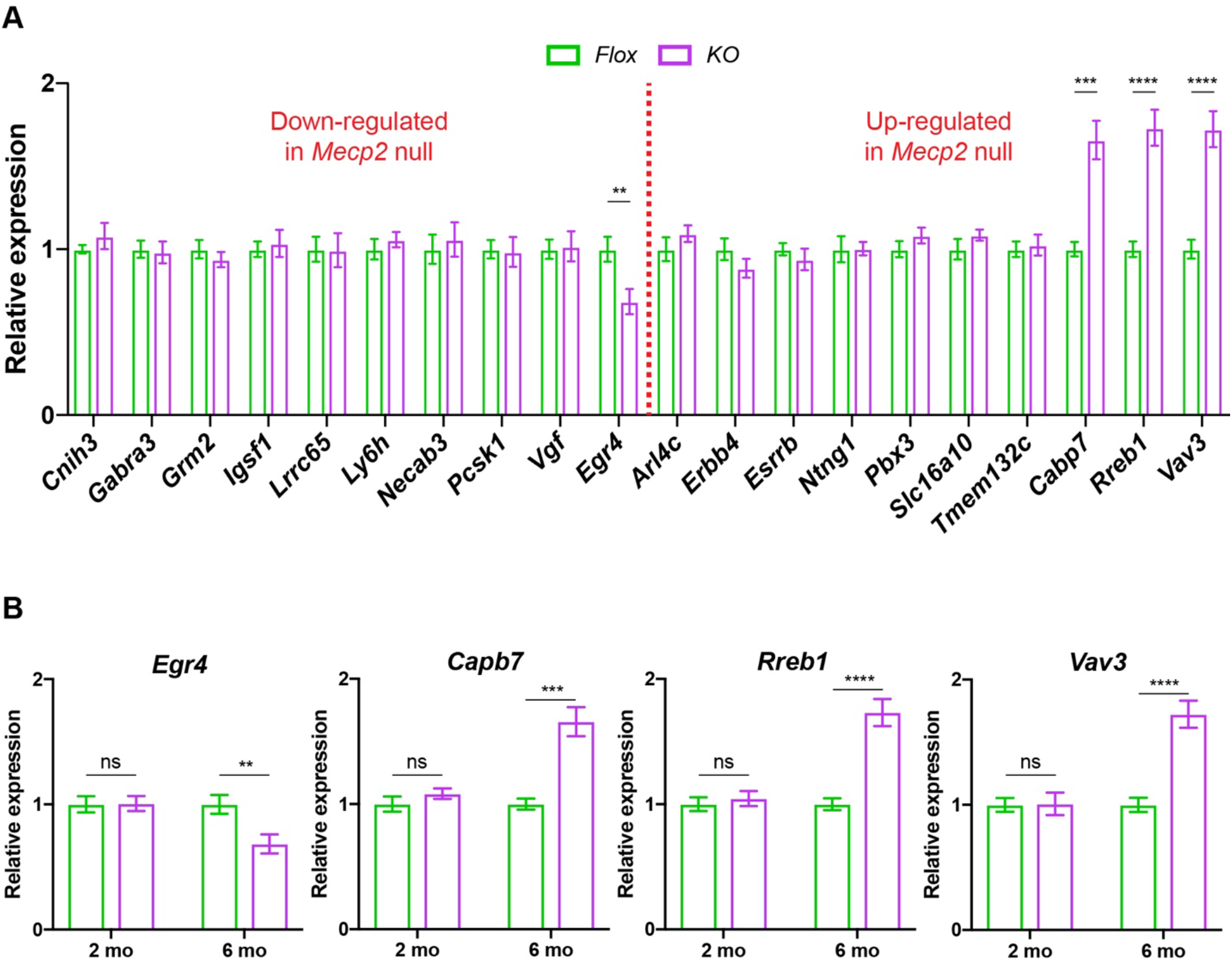
*KO* mice exhibit cerebellar gene expression changes. (**A**) Genes that are up- and down-regulated in global *Mecp2* null mice were analyzed by RT-qPCR in 6-month-old *Flox* and *KO* mice. (**B**) Expression levels of the misregulated genes in symptomatic 6-month-old *KO* mice were analyzed in asymptomatic 2-month-old *KO* mice. N = 6 biologically independent mice per group. Data are presented as mean ± s.e.m. ns(p>0.05), **(p<0.01), ***(p<0.001), ****(p<0.0001).

## DISCUSSION

Our study revealed three important features of cerebellar dysfunction that occur following the loss of *Mecp2*. First, we did not observe non-motor phenotypes in cerebellar *KO* mice even though the cerebellum is implicated in non-motor behaviors such as social interaction and cognition (Mauk et al., 2000; Tsai et al., 2012; Klein et al., 2016). This suggests that non-motor phenotypes in Rett syndrome are likely not caused by cerebellar dysfunction. Second, we did not observe motor deficits in any of the cell type-specific knockout mice. The same phenomenon is seen for the sensorimotor gating deficits of *Mecp2* null mice, which are present in mice lacking *Mecp2* in all inhibitory neurons but not from individual subtypes of inhibitory neurons (Chao et al., 2010; Ito-Ishida et al., 2015). This suggests that the cerebellar-related motor deficits are the result of combined dysfunction in the entire circuit rather than dysfunction in a single cell type. Finally, the behavioral deficits in cerebellar *KO* mice were milder than mice lacking *Mecp2* in the forebrain and basal ganglia. In cerebellar *KO* mice, the phenotypes were restricted to motor learning and appeared in six-month-old mice, whereas the motor deficits in mice lacking *Mecp2* in the forebrain and basal ganglia are more profound and arise in two-month-old mice (Chen et al., 2001; Gemelli et al., 2006; Su et al., 2015). Thus, the motor symptoms of Rett syndrome arise from a combination of cerebellar, forebrain, and basal ganglia dysfunction.

Although the synaptic and transcriptional changes in cerebellar KO mice were mild compared to those observed in other *Mecp2* knockout models (Chao et al., 2010, Meng et al., 2016; Raman et al., 2018), they may still contribute to the behavioral deficits since similar observations are observed in other mouse models of cerebellar dysfunction. For example, the loss of the ⍺3 isoform of the Na+/K+ pump in mice causes motor incoordination and dystonia (Calderon et al., 2011). In these mice, the mean firing rate of their Purkinje cells is normal, while the firing pattern is more irregular, similar to what we observed in cerebellar *KO* mice (Fremont et al., 2016). Just like cerebellar *KO* mice, the Purkinje cells of *Car8^wdl^* mice also have more irregular simple spike activity, which ultimately contribute to motor incoordination on the rotarod (White et al., 2016). Clearly, abnormal firing patterns of Purkinje cells lead to cerebellar dysfunction and motor phenotypes, which is likely occurring in cerebellar *KO* mice. At the transcriptional level, the misregulated genes in cerebellar *KO* mice play important roles in learning, neuronal function, and cerebellar function (Quevedo et al., 2010; Ludwig et al., 2011; Farley et al., 2018, Fairless et al., 2019).

A unique and interesting finding was the improvement in motor learning after additional training in cerebellar *KO* mice. To our knowledge, this phenomenon is not observed in other *Mecp2* knockout mice (Li and Pozzo-Miller, 2012; Lombardi et al., 2015). However, a related effect is seen in germline *Mecp2* null mice in which their memory deficits are rescued with forniceal deep brain stimulation (DBS) (Hao et al., 2015; Lu et al., 2016). The effects of DBS on brain circuitry share similarities to that of repetitive activation during training (De Zeeuw and Ten Brinke, 2015; Herrington et al., 2016; Lu et al., 2016; Langille and Brown, 2018). In cerebellar *KO* mice, it is possible that activation of the cerebellar circuitry during training might improve their motor phenotypes by improving synaptic function in a manner similar to the proposed mechanism of DBS. This also raises the possibility that repetitive circuit activation via training could improve other behavioral deficits in *Mecp2* knockout mice.

Taken together, our results reveal that cerebellar dysfunction contributes to motor deficits following the loss of MeCP2. Interestingly, the behavioral, synaptic, and transcriptional deficits in cerebellar *KO* mice were relatively mild compared to other *Mecp2* knockout models. So, even though MeCP2 is broadly expressed throughout the brain, neuronal cell types between brain regions, and even those within a brain region, respond differently to the loss of MeCP2. As future studies seek to better define the function of MeCP2 and use this knowledge to design effective therapies, it is important to keep in mind that the functional consequences of MeCP2 loss are context specific.

## MATERIALS AND METHODS

### Animals

Mice were maintained on a C57B/6J background on a 12 hr light: 12 hr dark cycle with standard mouse chow and water *ad libitum*. Mice were group housed up to five mice per cage. All behavioral experiments were performed during the light cycle at the same time of day. The following Cre-expressing mice were used for breeding: *En1-Cre* (*En1^tm2(cre)Wrst^*/J), *Atoh1-Cre* (B6.Cg-Tg(Atoh1-cre)1Bfri/J), *L7-Cre* (B6.129-Tg(Pcp2-cre)2Mpin/J), and *Ptf1a-Cre* (*Ptf1a^tm1(cre)Hnak^*/RschJ). Cre-expressing male mice were bred to *Mecp2^flox/+^* female mice to generate experimental groups (Chen et al., 2001). Mice were obtained from the Jackson Laboratories and maintained by breeding mice to wild-type C57B/6J mice. Only male offspring were used for experiments. Behavioral, histological, and electrophysiological analyses were performed blind to genotypes. The Baylor College of Medicine Institutional Animal Care and Use Committee approved all research and animal care procedures.

### Behavioral assays

For each test, mice were habituated in the room for 30 min. A light intensity of 150 lx and 60 dB background white noise was presented during habituation and testing. All assays were performed at the same time of day.

### Rotarod

Mice were placed on an accelerating rotarod apparatus (Ugo Basile) while the cylinder increased from 5 r.p.m to 40 r.p.m over a 5 min period. Latency to fall was measured when the mouse fell off the apparatus or rode the cylinder for two consecutive revolutions without regaining control. The trials consisted of four attempts with a 30 min rest after each trial. Data are represented as mean ± s.e.m. and analyzed by two-way ANOVA with Tukey’s multiple comparisons test.

### Eyeblink Conditioning

Eyeblink conditioning was performed as previously described (Heiney et al., 2014). Briefly, animals were anesthetized with isoflurane (1.5-2% by volume in O2, SuriVet). A midline incision was made to expose the skull and two small screws were placed on either side of the midline at bregma. A thin stainless steel headplate was placed on the skull such that the screws fit in the hole in the headplate. The plate was adhered to the skull using Metabond cement (Parkell). After 5 days of recovery, mice were habituated to the head restraint in the testing chamber for 2 days prior to training for 1 hr. Each training session consisted of 100 trials of the conditioned stimulus (CS, blue LED light) paired with the unconditioned stimulus (US, 20-30 psi periocular air puff). The interstimulus interval was 200 ms with an intertrial interval of at least 10 sec. The pressure of the periocular air puff was set for each mouse to elicit a full reflexive blink. The conditioned response (CR, eyelid closure) was monitored using infrared illumination and a high-speed camera (Allied Vision) combined with MATLAB using a custom-written software and acquisition toolbox. Mice were trained for 12 days after habituation. Eyelid traces were normalized to the full blink range. Trials were considered to contain a conditioned response if the eyelid closure exceeded 5% from the baseline mean within the CS-US interval. The CR probability was quantified as the number of conditioned responses divided by the total number of trials for each day. The CR amplitude was quantified as the maximum eyelid position within the CS-UR interval relative to the trial baseline position for each. Data are represented as mean ± s.e.m. and analyzed by two-way ANOVA with Tukey’s multiple comparisons test.

### Open Field Assay

Mice were placed in a clear, open Plexiglas box (40 x 40 x 30 cm, Stoelting) with an overhead camera and photo beams to record horizontal and vertical movements. Activity was measured over 10 min and quantified using ANY-maze (Stoelting). Data are represented as mean ± s.e.m. and analyzed by one-way ANOVA with Tukey’s multiple comparisons test.

### Parallel Rod Footslip

Mice were placed into the center of a wire grid laid in an open-field chamber (Accuscan) for 10 min. The number of footslips through the wire grid was recorded and analyzed using ANY-maze (Stoelting). The number of footslips was normalized to the total distance traveled. Data are represented as mean ± s.e.m. and analyzed by one-way ANOVA with Tukey’s multiple comparisons test.

### Grip Strength

Mice were held by the tail and allowed to grasp the bar of a grip strength meter (Chatillon-Ametek) with both forepaws. The mouse was pulled away from the bar until it released from the bar. The maximum force generated was averaged over three trials and normalized to the weight of the mouse. Data are represented as mean ± s.e.m. and analyzed by one-way ANOVA with Tukey’s multiple comparisons test.

### Wire Hang Time

Mice were allowed to grasp the middle of a 3 mm plastic coated wire suspended 6 in above a plastic-covered foam pad. The plastic wire was inverted for a maximum of 180 sec or until the mouse fell off. Data are represented as mean ± s.e.m. and analyzed by one-way ANOVA with Tukey’s multiple comparisons test.

### 3-Chamber Interaction

During the habituation phase, mice were placed in the middle of the three-chamber apparatus (Ugo Basile) containing two empty barred cages in the right and left chambers for 10 min. During the social interaction phase, an age-matched C57BL/6J wild-type male mouse was placed in one cage and a black Lego block of similar size was placed in the other. Partner mice were habituated to the chamber for 1 hr per day for two consecutive days before testing. The test mouse was returned to the middle zone and allowed to explore the chamber for 10 min. Mouse movement was recorded and analyzed using ANY-maze (Stoelting). Data are represented as mean ± s.e.m. and analyzed by two-way ANOVA with Tukey’s multiple comparisons test.

### Fear Conditioning Assay

Mice were trained in a fear-conditioning chamber (Med Associates, Inc.) that delivers an electric shock paired with a tone. On the first day, mice were placed in a holding room and delivered to the testing room in a temporary cage. This device was located inside a soundproof box that contained a digital camera and loudspeaker. Each mouse was placed individually in the chamber and left undisturbed for 2 min. A tone (80 dB, 5 kHz, 30 s) coincided with a foot-shock (2 s, 0.7 mA) and was repeated after 1 min. The apparatus was cleaned with isopropanol. The mouse was returned to the temporary cage after an additional minute and returned the home-cage in the holding room. Fear memory was assessed after 1 day of training. To test contextual fear memory, mice were placed in the original environment without a tone or foot-shock for 5 min. Mice were returned to their home-cage in the holding room. To test cued fear memory, mice were returned to the testing room and placed in the chamber, which was modified to distinguish it from the original context. The chamber was made triangular with the addition of white panels, cleaned with 70% ethanol, and scented with a cup of vanilla extract under the floor. The mouse was allowed to explore the novel environment for 3 min, after which the original tone (80 dB, 5 kHz, 3 min) was presented. Mouse movement was recorded and analyzed using ANY-maze (Stoelting). Freezing was scored only if the animal was immobile for at least 1 s. Data are represented as mean ± s.e.m. and analyzed by one-way ANOVA with Tukey’s multiple comparisons test.

### Acoustic Startle and Pre-pulse Inhibition

Mice were placed in a Plexiglas tube and allowed to habituate for 5 min with a 70 dB background noise. The test sessions consisted of six trials of each sound stimuli lasting 20 ms: no stimulus, a 120 dB sound burst, or a 120 dB sound burst with a 74 dB, 78 dB, or 82 dB pre-pulse stimuli presented 100 ms before the startle stimulus. The maximum startle response was recorded and analyzed during the 65 ms period following the onset of the startle stimulus (SR-Lab). Pre-pulse inhibition was calculated as 1 -(startle response with pre-pulse stimulus/startle response only) x 100). Data are represented as mean ± s.e.m. and analyzed by two-way ANOVA with Tukey’s multiple comparisons test.

### Histology and Immunofluorescence Staining

For immunofluorescence, animals were transcardially perfused with 50 mL ice-cold 4% paraformaldehyde in 0.1 M PBS. Brains were dissected, post-fixed overnight at 4 °C, washed with 0.1 M PBS, and placed in 30% sucrose in 0.1 M PBS for 24-48 hrs until the brains sunk to the bottom of the vial. Brains were embedded in Tissue-Tek Optimum Cutting Temperature Compound (Cat#4583, Sakura) and stored at −80 °C until further use. 50 μm floating sections were cut using a cryostat (Cat# CM3050S, Leica) and collected in 0.1 M PBS. Sections were incubated in blocking solution (0.3% Triton X-100, 5% normal goat serum in 0.1 M PBS) for 1 hr at room temperature followed by primary antibody in blocking solution for 24 hr at 4 °C. The following primary antibodies were used: rabbit anti-MeCP2 (1:1,000, Cat# D4F3, Cell Signaling Technology), mouse anti-MeCP2 (1:500, Cat# ab50005, Abcam), mouse anti-NeuN (1:250, Cat# MAB377, Millipore Sigma), mouse anti-Calbinin-D28K (1:10,000, Cat# 300, Swant), rabbit anti-parvalbumin (1:1,000, Cat# PV27, Swant), mouse anti-Vglut1 (1:1,000, Cat# 135 302, Synaptic Systems), guinea pig anti-Vglut2 (1:1,000, Cat# 135 404, Synaptic Systems), guinea pig anti-Vgat (1:1,000, Cat# 131 005, Synaptic Systems). Sections were washed with PBS and incubated in secondary antibody for 2 hr at room temperature. The following secondary antibodies were used: goat anti-mouse IgG Alexa Fluor 488 (1:500, Cat# A-11001, Thermo Fisher), goat anti-guinea pig IgG Alexa Fluor 555 (1:500, Cat# A-21435, Thermo Fisher), and goat anti-rabbit IgG Alexa Fluor 647 (1:500, Cat# A-21244, Thermo Fisher). Sections were counterstained with 1 mM DAPI (DAPI; Cat# D1306) for 5 min, washed with PBS, and mounted on electrostatic Superfrost Plus microscope slides (Cat# 12-550-15, Fisher Scientific) with ProLong Gold Antifade mounting medium (Cat# P10144, Thermo Fisher). Slides were cured overnight at room temperature and stored at 4 °C prior to imaging.

For Golgi-Cox staining, the brains were removed from the skull and processed using the FD Rapid Golgi Stain Kit (Cat #PK401, FD Neurotechnologies). All steps were carried out according to the manufacturer’s instructions. The tissue was sectioned at 100 μm and transferred to electrostatic glass slides and mounted with ProLong Gold Antifade mounting medium (Cat# P10144, Thermo Fisher). For Nissl staining, 50 μm frozen sections were mounted onto electrostatic glass slides, dehydrated overnight in 1:1 ethanol:chlorofom, rehydrated in 100 % and 95 % ethanol, stained with 0.1 % cresyl violet, washed in 95 % and 100 % ethanol, cleared with xylene, and mounted with mounting medium. Slides were stored at room temperature prior to imaging.

### Imagining and Quantification

Brightfield images were captured with the AxioCam MRc5 camera (Zeiss) mounted on an Axio Imager (Zeiss). Confocal images were captured with the TCS SP8 microscope (Leica) using a 10X or 63X objective. Z-stack images were acquired at 10 μm steps for 10X images, or 0.5 μm steps for morphological analysis and 63X images. Laser settings were set above background levels based on the signal intensity of tissue stained only with the secondary antibody and kept consistent across samples in each experiment. Vglut1, Vglut2, and Vgat puncta were counted using the Analyze Particles function in ImageJ-Fiji (Schindelin et al., 2012). Neuronal structure reconstruction, Sholl analysis, and spine quantification was performed with Neuroludica 360 (MBF Bioscience).

### Purkinje Cell Electrophysiology

Single-unit extracellular recording was performed as previously described (Heiney et al., 2018). A 2-3 mm-diameter craniotomy was opened over the right side of the cerebellum (6.5 mm posterior and 2.0 mm lateral from bregma), and the dura was protected by a layer of Kwik-Sil (WPI). A custom 3D printed recording chamber and interlocking lid (NeuroNexus) was secured over the craniotomy with dental acrylic to provide additional protection. After 5 days of recovery, the mouse was fixed in place on a treadmill via a previously implanted headplate. Purkinje cell simple spikes (SSpk) and complex spikes (CSpk) were isolated using a tetrode (Thomas Recording, AN000968) acutely driven into the cerebellar cortex with microdrives mounted on a stereotactic frame (Narishige MMO-220A and SMM-100). We targeted Purkinje cells in lobule V located at the ventral part in the medial wall of the primary fissure, which includes the cerebellar microzone that supports eyeblink conditioning (Heiney et al., 2014; Ohmae & Medina, 2015). Targeting lobule V instead of widely sampling from the cerebellum allowed us to compare the statistical results of spike firing in Purkinje cells in the same cerebellar subregions across control and mutant mice while minimizing the effect of zebrin-derived difference on the statistical results, because Purkinje cells in the lobule V are largely zebrin negative (Zhou et al., 2014, Saša et al., 2016;). The final recording site in each mouse was marked by an electrical microlesion (0.01 mA, 60 seconds), and the other locations of recorded Purkinje cells were reconstructed based on the stereotaxic position of each recording location relative to the lesion site. Recording data were excluded from further analysis if Purkinje cells were not located in lobule V. Data were recorded and stimuli were delivered using an integrated Tucker-Davis Technologies and Matlab system (TDT RZ5, medusa, RPVdsEx) running custom code (github.com/blinklab/neuroblinks). SSpks and CSpks were sorted manually offline for the duration of recordings (mean ± SEM, 310 ± 61s). The different spike types were identified by their characteristic waveforms in Spike2, and recordings with poor SSpk and CSpk isolation were excluded from further analysis. Inter-spike intervals (ISIs) were calculated for consecutive simple spikes, and CV2 was computed by 2 x |ISI_n_ – ISI_n-1_| / (ISIn + ISI_n-1_) and averaged across all consecutive ISIs within neuron.

### Western Blotting

The cerebellum of cKO and control mice were rapidly dissected, flash frozen in liquid nitrogen, and stored at −80 °C until further use. The cerebellum was placed in a glass homogenizer with ice-cold lysis buffer (2% SDS, 100 mM Tris-HCl, pH 7.5, protease inhibitor, and phosphatase inhibitor). The cerebellum was homogenized with 10 strokes of pestle A and 10 strokes of pestle B. Samples were sonicated on the Bioruptor sonication device (Cat# B01060010, Diagendoe) for 30 sec ON and 30 sec OFF for 10 cycles. The homogenate was centrifuged at 10,000 x g for 10 min at 4 °C. Protein concentration from the supernatant was measured using a Pierce BCA Protein Assay Kit (Cat# 23225, ThermoFisher Scientific). Lysates were diluted to 1 μg/μl in an extraction buffer of 1 X NuPAGE LDS Sample Buffer (Cat# NP0007, ThermoFisher Scientific), 1 X NuPAGE Sample Reducing Agent (Cat# NP0004, ThermoFisher Scientific), and 1 X RIPA buffer (100 mM Tris pH 8.0, 300 mM NaCl, 0.2 % SDS, 1 % sodium deoxycholate, 2 % Nonidet P-40, 5 mM EDTA, protease inhibitor, and phosphatase inhibitor). Samples were then heated at 95 °C for 10 min. 10 μg of protein was loaded into a 15-well NuPAGE 4-12 % Bis-Tris gel (Cat# NP0336BOX, ThermoFisher) and run in MES buffer (50 mM MES, 50 mM Tris base, 0.1 % SDS, 1 mM EDTA pH 7.3) for 20 min at 200 V. Proteins were transferred onto Amersham Hybond 0.2 PVDF blotting membranes (Cat# 10600021, GE Healthcare Life Sciences) in Tris glycine buffer (25 mM Tris, 190 mM glycine, 20 % methanol). Membranes were rinsed in ddH2O and dried at room temp for 1 hr. Membranes were rehydrated in methanol, blocked for 1 hr at room temperature with Odyssey TBS Blocking Buffer (Cat# 927-50000, LI-COR Biosciences) and incubated in primary antibody diluted in blocking buffer overnight at 4 °C. The following primary antibodies were used: rabbit anti-MeCP2 (1:1,000, Cat# D4F3, Cell Signaling Technology), rabbit anti-Histone H3 (1:20,000, Cat# ab1791). The membranes were washed 3 times with TBS-T for 10 min each, then incubated in secondary diluted in blocking buffer for 2 hr at room temperature. The following secondary antibody was used: IRDye 680RD goat anti-rabbit IgG (Cat# 926-6807, LI-COR Biosciences). The membranes were washed 3 times with TBS-T for 10 min each and imaged on an Odyssey imager (LI-COR Biosciences). Relative intensities were quantified using ImageStudioLite (LI-COR Biosciences).

### RNA Isolation and RT-qPCR

The cerebellum of cKO and control mice were rapidly dissected, flash frozen in liquid nitrogen, and stored at −80°C until further use. RNA was harvested using the RNeasy Mini Kit (Cat# 74104, Qiagen). 500 ng of purified RNA was reverse transcribed using M-MLV Reverse Transcriptase (Cat# 2802501, ThermoFisher Scientific) and random hexameters primers (Cat# 48190011, ThermoFisher Scientific). Newly synthesized cDNA was diluted 1:20 (vol:vol) in pure H2O and analyzed by qPCR using PowerUp SYBR Green Master Mix (Cat# A25742, ThermoFisher Scientific) on a CFX96 Real-Time PCR Detection System (Cat# 1855195, Bio-Rad). The following primers were used: 5’-CGCTCATCTTTTTCGCTATCTGG-3’ and 5’-CTTGAAATCCGTTCTTAGCTCGT-3’ (*Cnih3*), 5’-ATGTGGCACTTTTATGTGACCA-3’ and 5’-CCCCAGGTTCTTGTCGTCTTG-3’ (*Gabar3*), 5’-GCTCCCACAGCTATCACCG-3’ and 5’-TCATAACGGGACTTGTCGCTC-3’ (*Grm2*), 5’-CTGCTCGGTGTCGCAACAT-3’ and 5’-GACATTGAGGTTCAGGAGGGC-3’ (*Igsf1*), 5’-CGTAACCAAGTAGTGGACTGC-3’ and 5’-TAGCATGTGAGATAGCCTGGC-3’ (*Lrrc65*), 5’-AGCCCACCGATACCGTTTG-3’ and 5’-AGAAAAAGTGCCGCTTAACGAA-3’ (*Ly6h*), 5’-GGAACCCACACATGAGGAGG-3’ and 5’-GCAGCACAAAGCACGATGAAA-3’ (*Necab3*), 5’-CGACGAGACTCCTGACGTG-3’ and 5’-GCACCTCGGGACCCAAATC-3’ (*Pcsk1*), 5’-AAGGATGACGGCGTACCAGA-3’ and 5’-TGCCTGCAACAGTACCGAG-3’ (*Vgf*), 5’-CGCGCAGTGACGAGAAGAA-3’ and 5’-GAGAGGCCCAGCGAGTAGA-3’ (*Egr4*), 5’-AGTCTCTGCACATCGTTATGC-3’ and 5’-GGTGTTGAAGCCGATAGTGGG-3’ (*Arl4c*), 5’-GTGCTATGGACCCTACGTTAGT-3’ and 5’-TCATTGAAGTTCATGCAGGCAA-3’ (*Erbb4*), 5’-GGACTCGCCGCCTATGTTC-3’ and 5’-CGTTAAGCATGTACTCGCATTTG-3’ (*Esrrb*), 5’-ACACACTGTACTAGGCCCTGA-3’ and 5’-CGTCCTGACTCGAATCTCCCA-3’ (*Ntng1*), 5’-ACCTCCCAAATTCTGGGGACA-3’ and 5’-ATCCACCTGTGACTGCACATT-3’ (*Pbx3*), 5’-AGGTGCTCTTCATGTGCATTG-3’ and 5’-TGGAGGTAGACCTTCTTCACAC-3’ (*Slc16a10*), 5’-TCAGAGCCGAGACTGCATTCT-3’ and 5’-GCCCATAGCTGACGTTTAATACC-3’ (*Tmem132c*), 5’-ATGTACCGGGGCATCTACAC-3’ and 5’-GGGCATGTAACCCAGGGAG-3’ (*Cabp7*), 5’-AGTAATGAGCGTAGCGAGTGT-3’ and 5’-GGTCCTGGAGGTTTCATGGG-3’ (*Rreb1*), 5’-CCTGCTGCGATACCTTTGGAA-3’ and 5’-GTGTTCGGGATAGCCGAGATA-3’ (*Vav3*).

### Statistical Analysis

Data are displayed as mean ±s.e.m. and the significance threshold was set at ⍺=0.05 (ns, *p<0.05, **p<0.01, ***p<0.001, ****p<0.0001). Sample sizes were determined based on prior statistics and data characterizing the phenotypes of MeCP2 mutant mice (Chao et al., 2010; Ito-Ishida et al., 2015; Meng et al., 2016). Statistical analysis was performed using Prism (GraphPad). Data was analyzed with the experimenters blinded to genotype.

## ACKNOWLEDGMENTS

This project was funded by the National Institutes of Health (R01NS057819 to H.Y.Z.; F30HD09787 to N.P.A.; R01MH093727, R01NS112917, and RF1MH114269 to J.F.M.; F31NS103427 to O.A.K.; R01NS089664 and R01NS100874 to R.V.S.), the Howard Hughes Medical Institute (H.Y.Z.), and the Baylor College of Medicine Intellectual and Developmental Disabilities Research Center (NIH 5U54HD083092). We thank all members of the Zoghbi and Medina labs for their critical review of the manuscript. Figure diagrams were created with BioRender.com.

---

**Figure 2—figure supplement 1.**
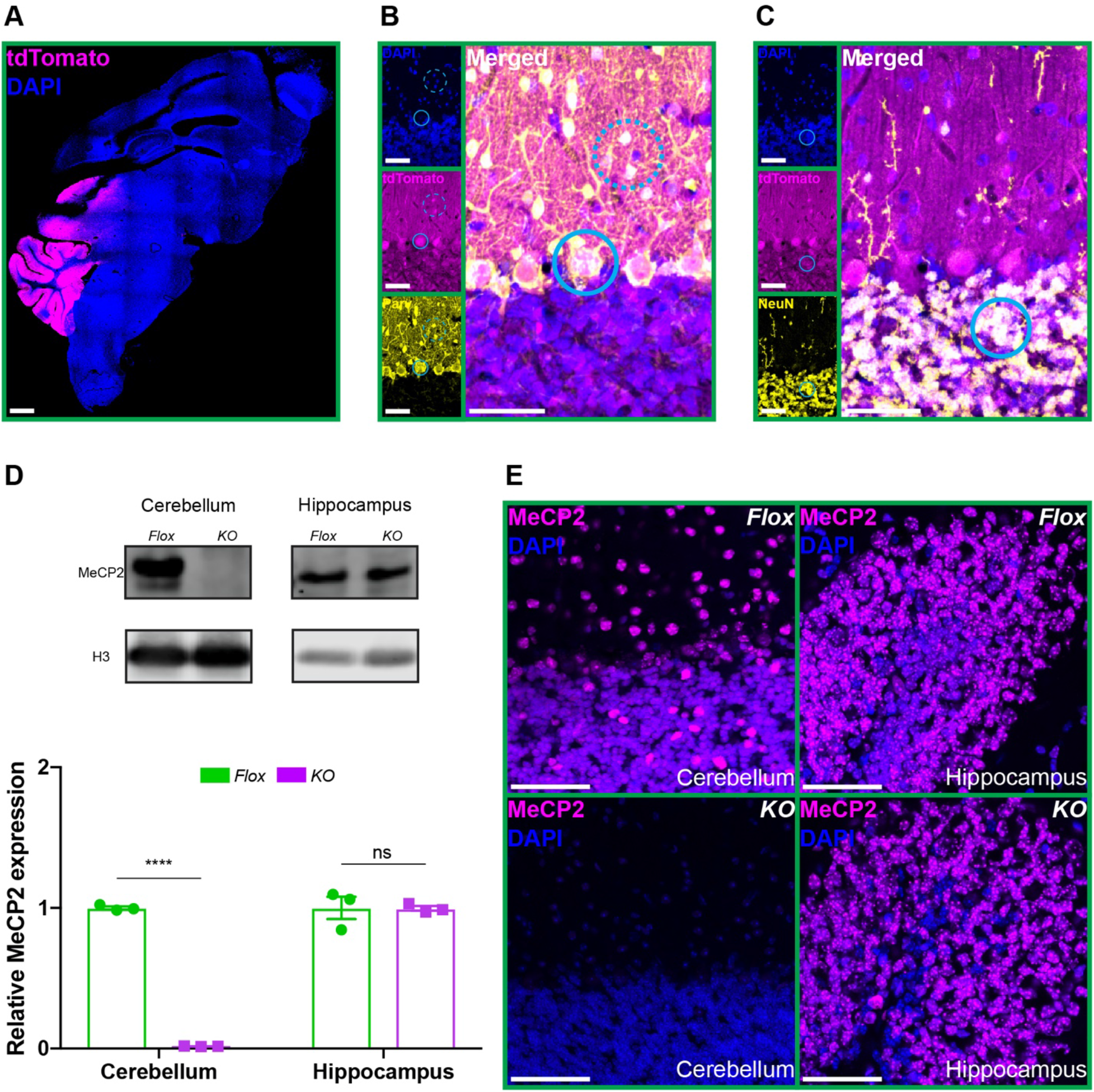
Deletion of *Mecp2* from the entire cerebellum. (**A**) *En1-Cre* expression was determined by the pattern of tdTomato (magenta) in *En1-Cre^+/^;Rosa26^tdTomato^* reporter mice. Scale bar, 1 mm. (**B**) Immunostaining showing co-expression of tdTomato (magenta) with Parvalbumin (yellow), a marker of Purkinje cells (solid cyan circle) and molecular layer inhibitory neurons (dashed cyan circle). Scale bar, 25 μm. (**C**) Immunostaining showing co-expression of tdTomato (magenta) with NeuN (yellow), a maker of granule neurons (solid cyan circle). Scale bar, 25 μm. (**D**) Quantification of MeCP2 protein levels normalized to Histone H3 in the cerebellum and hippocampus of *Flox* and *KO* mice. (**E**) Immunostaining of MeCP2 (magenta) in the cerebellum and hippocampus of *Flox* and *KO* mice. Scale bar, 50 μm. N = 3 biologically independent mice per group. Data are presented as mean ± s.e.m. ns(p>0.05), ****(p<0.0001).

**Figure 2—figure supplement 2.**
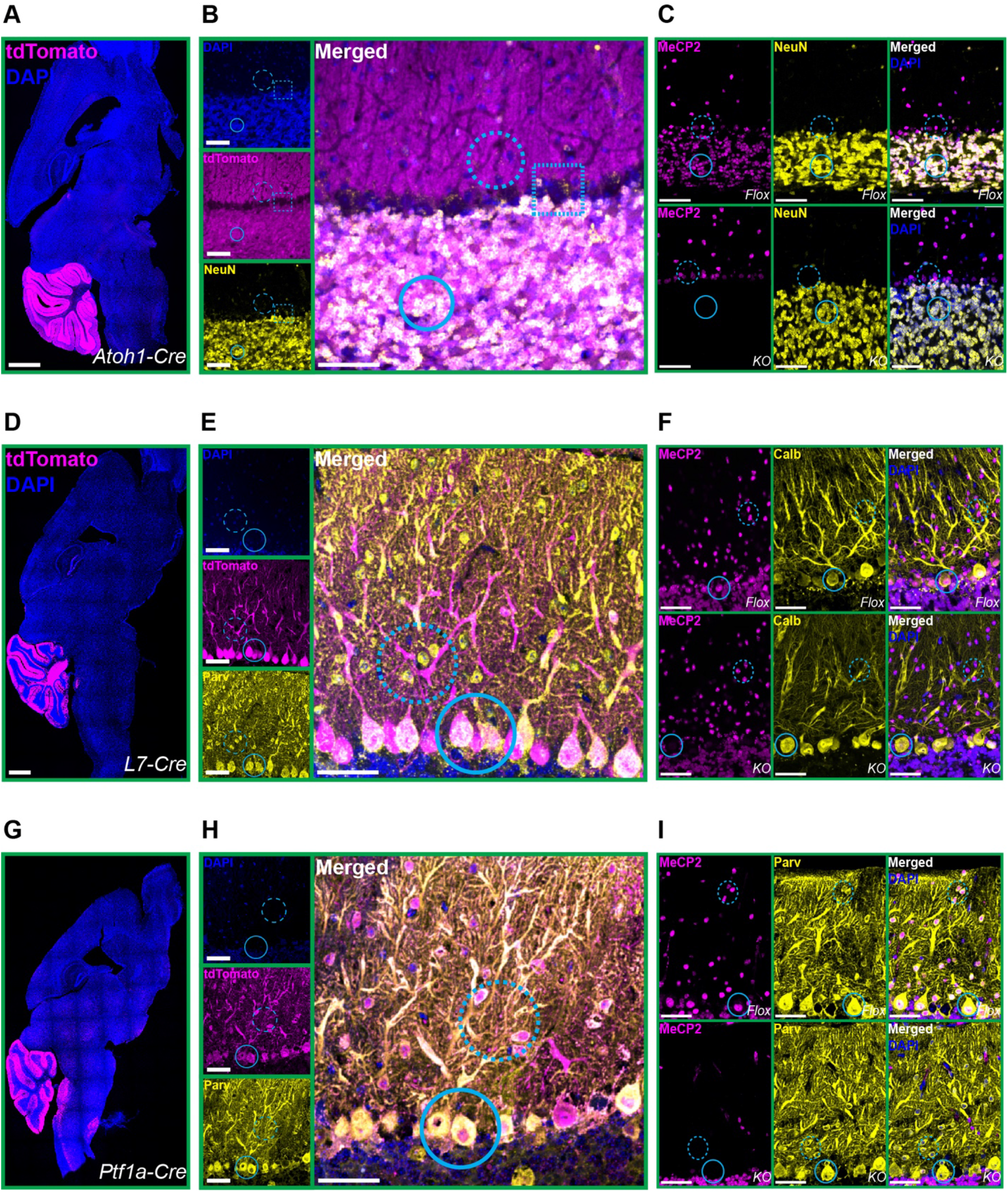
*Cre* expression and *Mecp2* deletion in individual cerebellar neuron types. (**A**) *Atoh1-Cre* expression was determined by the pattern of tdTomato (magenta) in *Atoh1-Cre^+/−^;Rosa26^tdTomato^* reporter mice. Scale bar, 1 mm. (**B**) Immunostaining in *Atoh1-Cre^+/−^;Rosa26^tdTomato^* reporter mice showing co-expression of tdTomato (magenta) with NeuN (yellow), a marker of granule cells (solid cyan circle), but not in cells of the Purkinje layer (dashed cyan square) or molecular layer (dashed cyan circle). Scale bar, 25 μm. (**C**) Immunostaining of MeCP2 (magenta) and NeuN (yellow) in the cerebellum showing the absence of MeCP2 in granule neurons of *KO* mice (solid cyan circle) but not in Purkinje layer neurons (dashed cyan circle). Scale bar, 25 μm. (**D**) *L7-Cre* expression was determined by the pattern of tdTomato (magenta) in *L7-Cre;Rosa26^tdTomato^* reporter mice. Scale bar, 1 mm. (**E**) Immunostaining in *L7-Cre;Rosa26*^tdTomato^reporter mice showing co-expression of tdTomato (magenta) with Parvalbumin (yellow) in the Purkinje cell layer (solid cyan circle) but not the molecular layer (dashed cyan circle). Scale bar, 25 μm. (**F**) Immunostaining of MeCP2 (magenta) and Calbindin (yellow) in the cerebellum showing the absence of MeCP2 in Purkinje cells of *KO* mice (solid cyan circle) but not in molecular layer neurons (dashed cyan circle). Scale bar, 25 μm. (**G**) *Ptf1a-Cre* expression was determined by the pattern of tdTomato (magenta) in *Ptf1a-Cre;Rosa26^tdTomato^* reporter mice. Scale bar, 1 mm. (**H**) Immunostaining in *Ptf1a-Cre;Rosa26^tdTomato^* reporter mice showing co-expression of tdTomato (magenta) with Parvalbumin (yellow) in the molecular cell layer (solid cyan circle). Scale bar, 25 μm. (**F**) Immunostaining of MeCP2 (magenta) and Parvalbumin (yellow) in the cerebellum showing the absence of MeCP2 in Purkinje cells (solid cyan circle) and molecular layer neurons of *KO* mice (dashed cyan circle). Scale bar, 25 μm.

**Figure 2—figure supplement 3.**
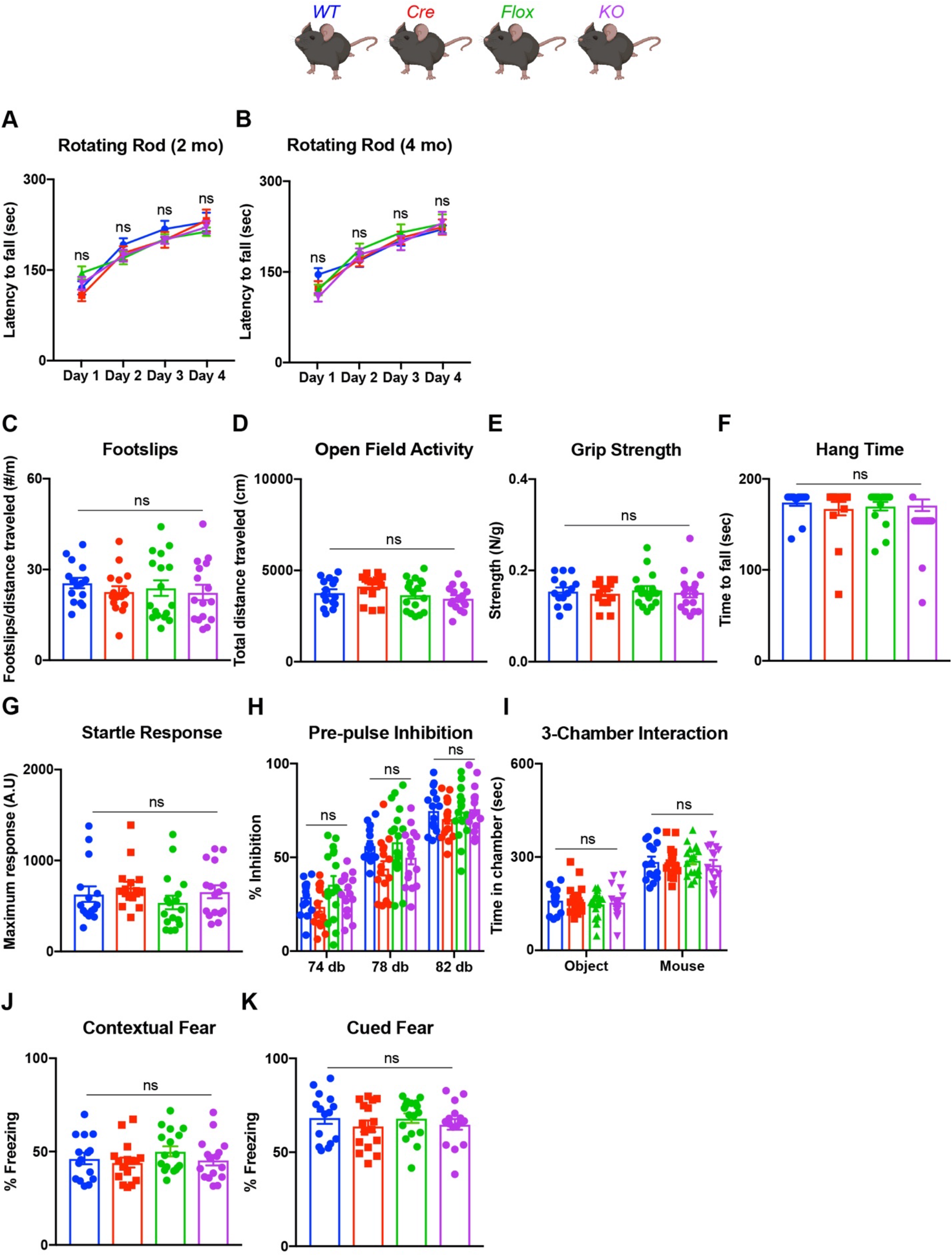
Deleting *Mecp2* from the entire cerebellum does not cause other behavioral abnormalities in 6-month-old mice. (**A**) Latency to fall on the rotarod in 2-month-old mice. (**B**) Latency to fall on the rotarod in 4-month-old mice. (**C**) Footslip count on the parallel rod assay normalized to total distance traveled. (**D**) Total distance traveled in the open field assay. (**E**) Grip strength normalized to body weight. (**F**) Hang time on an inverted wire grid. (**G**) Maximum acoustic startle response to a 120 dB stimulus. (**H**) Pre-pulse inhibition at 74, 78, and 82 dB pre-pulses. (**I**) Interaction time in the 3-chamber social interaction assay between a novel mouse or object. (**J**) Time spent freezing during contextual memory recall. (**K**) Time spent freezing during cued memory recall. N = 10-17 biologically independent mice per group. Data are presented as mean ± s.e.m. ns(p>0.05).

